# Metabolic analysis of a bacterial synthetic community from maize roots provides new mechanistic insights into microbiome stability

**DOI:** 10.1101/2021.11.28.470254

**Authors:** Jenna Krumbach, Patrizia Kroll, Vera Wewer, Sabine Metzger, Till Ischebeck, Richard P. Jacoby

## Abstract

Stability is a desirable property for agricultural microbiomes, but there is a poor understanding of the mechanisms that mediate microbial community stability. Recently, a representative bacterial synthetic community from maize roots has been proposed as a model system to study microbiome stability (Niu 2017, PNAS, 114:E2450). This SynCom assembles stably when all seven members are present, but community diversity collapses without the keystone E. cloacae strain. The aim of this study was to assess the role of metabolites for the stability of this SynCom, by defining the metabolic niches occupied by each strain, as well as their cross-feeding phenotypes and B-vitamin dependencies. We show that the individual member strains occupy complementary metabolic niches, measured by the depletion of distinct metabolites in exometabolomic experiments, as well as contrasting growth phenotypes on diverse carbon substrates. Minimal medium experiments show that the established seven-member community comprises a mixture of prototrophic and auxotrophic strains. Correspondingly, experimental cross-feeding phenotypes showed that spent media harvested from the prototrophic strains can sustain growth of two auxotrophs. We suggest that the metabolic mechanisms exhibited by this SynCom could serve as design principles to inform the rational assembly of stable plant-associated microbial communities.

## Introduction

A deeper understanding of microbiome stability is a major research goal in the field of microbiome science and technology (1). Potentially, stability criteria could help to predict how different microbiomes respond to disruption, such as environmental change or biotic invasion (2). Furthermore, a mechanistic understanding of microbiome stability could help to rationally formulate bio-inoculants in agriculture and medicine, via combining cooperative strains into stable synthetic communities that will persist in the target environment (3).

Conceptually, metabolism is positioned as a central factor mediating microbiome stability, underpinned via two major processes: 1) Niche differentiation, and 2) Cross-feeding (4,5). The ecological principle of niche differentiation stipulates that organisms which coexist in the same habitat must avoid competition by consuming different resources (6). Although theoretical studies predict that niche differentiation plays a major role in facilitating the diversity of microbiome composition, there are still significant knowledge gaps regarding the specific metabolites consumed by individual strains (7). Meanwhile, cross-feeding is also proposed to facilitate microbial diversity, because the metabolites released by one strain could provide a set of new nutrient niches to support the coexistence of other strains, and furthermore because the release of exogenous vitamins and amino acids would also support microbial diversity by nourishing auxotrophic strains (8). However, there is a relatively poor mechanistic knowledge of which particular microbial strains engage in cross-feeding, and what specific metabolites are shared (9).

The laboratory study of metabolic interactions in microbial communities has been hampered by the lack of experimentally tractable systems (10). However, a recently characterised seven-member bacterial synthetic community from maize roots is emerging as a model system for plant microbiome research (11). Notably, this SynCom assembles stably when all seven members are present, but the removal of strain Ecl results in a collapse of microbial community diversity. This observation is reminiscent of the keystone effect in macroecology, whereby one particular species plays a disproportionately large role in facilitating ecosystem diversity (12).

The objective of this study was to describe the metabolic characteristics of this previously established seven-member bacterial SynCom, by exploring the concepts of niche differentiation and cross-feeding. We conducted experiments to: 1) Assess the metabolic niche of each strain, 2) Describe cross-feeding interactions between donor and recipient strains, and 3) Define the B-vitamin responses of SynCom members. These measurements help to interpret the data of Niu et al (11), by providing mechanistic explanations for why certain strains play a disproportionate role in structuring community assembly.

## Methods

### Exometabolomics of bacterial cultivated on maize root extract

Microbial exometabolomics on maize root extracts was conducted according to the method adapted from Jacoby et al (13). Maize seeds (v. Sunrise) were purchased from Agri-Saaten Ltd, and sterilised in 70% ethanol for 15 min, followed by 5% NaHClO_4_ for 15 min, rinsed five times with sterile water, then incubated in sterile water at 50° C for 10 min. Individual seeds were then placed in Petri dishes with 10 mL of sterile water and incubated in the dark for 4 days at 30° C. Next, germinated seedlings were transferred into a sterile hydroponic growth system, enclosed in transparent plastic boxes with HEPA-filters for gas exchange (Sac O_2_ Ltd), with roots supported by 3 mm glass beads, and growth medium containing 0.5× Hoagland’s solution. Plants were placed in a growth chamber with 23/18° C day/night temperature, 16 h daylength, and 150 μmolm^-1^s^-1^ light intensity. Plants were cultivated for seven days, with growth medium changed once. At harvest, roots were separated from shoots, washed three times in sterile 10 mM MgCl_2_, then root tissue was snap-frozen in liquid N_2_.

For metabolite extraction, frozen maize root tissue was ground to a fine powder using a mortar and pestle and liquid nitrogen. Next, 200 mg of tissue powder was placed into a 1.5 mL tube, and incubated with 1 mL of 90% MeOH at 60° C for 30 min with 1500 rpm shaking. Tubes were centrifuged at 10,000× g for 10 min, then 800 μl of supernatant was transferred to a new tube and dried down in a vacuum centrifuge overnight. Next, dried metabolites were dissolved in water and filter-sterilised (0.22 μm pore size). Total organic carbon (TOC) concentration was measured using a Dimatoc 2000 (Dimatec).

For bacterial pre-culture, strains were streaked from glycerol stocks onto TSA plates (0.5× TSB, 1.2% agar) and incubated at 28° C for 1-2 days. Individual colonies were picked and transferred into 4 mL of 0.5× TSB at 28° C with 200 rpm shaking for 1-2 days. Cells were harvested by centrifuging 900 mL of culture at 5,000× g for 5 min at RT. Cell pellets were then washed twice with 900 mL of sterile 10 mM MgCl_2_, and resuspended at a final OD600 of 1 in sterile 10 mM MgCl_2_. The C7 mixture of all seven strains was prepared by equally combining washed bacterial cells at OD=1.

Next, bacteria were cultivated on an M9 growth medium where maize root extracts were the sole carbon source. The medium contained 24 mM Na_2_HPO_4_, 20 mM NH_4_Cl, 11 mM KH_2_PO_4_, 4 mM NaCl, 1 mM MgSO_4_, 100 μM CaCl_2_, 50 μM Fe-EDTA, 50 μM H_3_BO_3_, 10 μM MnCl_2_, 1.75 μM ZnCl_2_, 1 uM KI, 800 nM Na_2_MoO_4_, 500 nM CuCl_2_, and 100 nM CoCl. To this, maize root extracts were added to a final concentration of 720 μg C per mL. Growth assays were conducted in a 48-well plate, by adding 20 μL of resuspended bacterial pellet into 380 μL of medium, and incubating the plate in a plate reader (Tecan Infinite Pro 100) at 28° C for 24 h, with shaking (3 min continuous orbital shaking followed by 7 min stationary, shaking amplitude 3 mm). A negative control (no bacteria) was also prepared by adding 20 uL of sterile 10 mM MgCl_2_ to the growth medium, and was incubated side-by-side with the bacterial cultivations. Spent media were harvested by centrifuging cultures at 10,000x g for 3 min, then filter-sterilising the derived culture supernatants using 0.22 μm spin filters.

For untargeted LC-MS analysis of culture supernatants, 5 μl of sample was loaded onto a C18 column (XSelect HSS T3, 2.5 μm particle size, 10-nm pore size, 150 mm length by 3.0 mm width; Waters), using an HPLC (Dionex Ultimate 3000, Thermo Scientific). Buffer A was 0.1% FA in water, buffer B was 0.1%FA in methanol, and flow rate was 500 μl/min. Samples were eluted using the following gradient: hold at 1% B between 0 to 1 min, linear increase to 40% B until 11 min, linear increase to 99% B until 15 min, hold at 99% B until 16 min, linear decrease to 1% B until 17 min, and finally, hold at 1% B until 20 min. MS was conducted using a Q-TOF MS (maXis 4G; Bruker Daltonics), following ESI. The MS was operated in both positive and negative-ion modes, using N_2_ as drying gas at a flow rate of 8 litres/min, dry heater set to 220° C, nebulizer pressure of 1.8 bar, capillary voltage of 4,500 V, and collision radio frequency voltage of 370 V (corresponding to a mass range of 50 to 1,300 m/z). Scan rate was 1 Hz.

To process the untargeted LC-MS data, raw .D files were centroided and converted to .MZMXL using the MSConvert program (14). Files were then uploaded to XCMS online (15) and were aligned using the following parameters: m/z tolerance of 15 ppm, minimum peak width of 10 s, maximum peak width of 60 s, signal/noise threshold of 10, overlapping peaks split when m/z difference was greater than 0.01 m/z, features only considered if they occurred in at least five consecutive scans with an intensity greater than 5,000. Also, features were only considered if they occurred in at least two of three replicates from any sample group. The CAMERA algorithm was implemented to detect isotopes and adducts. Following export of files from XCMS online, data from both positive and negative-ion modes were combined, and filtered to remove isotopic peaks, and also filtered to only include MS features with RT between 1 to 16 min.

The statistical analysis of the LC-MS exometabolomic data had two aims: 1) To detect maize metabolite ions that were depleted from the medium following bacterial growth, and 2) To detect microbe-derived metabolite ions that were enriched in the medium following bacterial growth. Both strategies involved comparing the metabolomic profiles of the inoculated samples versus the uninoculated negative control. For a metabolite ion to be categorised as depleted, thresholds were: signal intensity < 50% versus sterile control, p-value < 0.05. For a metabolite ion to be categorised as enriched, thresholds were signal intensity > 200% versus sterile control, p-value < 0.05.

To assign candidate IDs to the metabolite ions of interest, m/z values were searched using CEU Mass Mediator (16) against the Metlin database (17) with 10 ppm tolerance, for various common adducts (positive mode: M+H, M+Na, M+K; negative mode: M-H, M+Cl, M+FA). In cases where multiple database compounds matched to the same m/z value, the compound with the lowest Metlin database identifier was chosen. These candidate metabolite IDs were then grouped into chemical categories using the ClassyFire database (18).

For GC-MS quantification of bacterial primary metabolite depletion from maize root extracts, 50 μl of spent or fresh medium was pipetted into 500 μl of Methanol/Chloroform/Water (5:2:1 (v/v/v)), and 200 of *allo*-inositol (5 μg/ml) was introduced into each sample as an internal standard. Next, 100 μL of the polar fraction was dried under a stream of N_2_ gas, and derivatised with 15 μl methoxyamine hydrochloride in pyridine (30 mg/ml) and 30 μl N-Methyl-N-(trimethylsilyl) trifluoroacetamide (MSTFA) (19). The samples were analysed on an Agilent 5977N mass selective detector connected to an Agilent 7890B gas chromatograph, as previously described (20). Primary metabolites were quantified according to the intensity of reporter ions previously obtained for pure reference compounds, normalised to the intensity of the *allo*-inositol internal standard. Bacterial depletion of primary metabolites was measured by comparing the normalised metabolite abundance in the inoculated samples versus the sterile controls.

### Phenotype microarray

Phenotype microarrays were conducted using BIOLOG EcoPlate according to the manufacturer’s instructions. The seven bacterial strains were pre-cultured, harvested and washed as described above, then diluted to an OD value of 0.1 in sterile 10 mM MgCl_2_. At this point, strains were mixed equally mixed together to compose either C7 communities or C6 drop-outs, and 150 μL of bacterial suspension was inoculated into each well of the 96 well-plate. Plates were incubated at 28° C for two days, and A590 was measured using a Tecan M100 Infinite Pro.

For analysis of phenotype microarray data, A590 values from all substrate-containing wells were compared to a negative control (no substrate) via Student’s t-test, and growth was considered positive if p-value < 0.05 and A590 > 0.1. In our hands, two substrates (2-hydroxy-benzoic acid and alpha-D-lactose) were not utilised by any bacterial inoculation and were therefore excluded from the analysis.

### Minimal medium growth assays

For assays of bacterial growth on single carbon substrates, bacteria were pre-cultured, harvested and washed as described above. Medium composition was the same as the M9 minimal medium described above, where all individual carbon sources were included at a concentration of 720 μg C per mL. Growth assays were conducted in a 96-well plate, where each well contained 95 μL of medium, inoculated with 5 μL of bacterial suspension, and cultivated in a plate reader (Tecan M100 Infinite Pro) using the same program as described above, except that data was collected over two days. Growth curves were quantitatively analysed using the Growthcurver R package (21).

### Bacterial cross-feeding assays

To analyse pairwise cross-feeding phenotypes across different carbon sources, the assay began by pre-culturing the three donor strains which could successfully grow on minimal medium (Ecl, Hfr and Ppu). This was conducted by picking colonies from TSA plates and transferring them into 4 mL of minimal medium, formulated as described above with either glucose, malate or alanine as sole carbon source at 720 μg C per mL, then incubating at 28° C with 200 rpm shaking for 2 days. The derived culture supernatants were harvested by centrifuging at 10,000x g for 3 min, then filter-sterilising the supernatants using 0.22 μm filters. In parallel, all seven recipient strains were pre-cultured on 0.5× TSB, harvested and washed as described above. In a 96-well plate, 5 μL of bacterial suspensions (OD=1) from the recipient strains were inoculated into 95 μL culture supernatants harvested from the donor strains, alongside a negative control of sterile 10 mM MgCl_2_ that was included to check for sterility of culture supernatants. Growth assays and quantitative analysis were performed as described above.

### Vitamin response assays

Several methodological strategies were used to dissect the vitamin responses of these strains. First, the responses of all seven strains to a mixture of B-vitamins was undertaken, whereby bacteria were pre-cultured, harvested and washed as described previously, and inoculated into a minimal medium containing either all eight B-vitamins or no vitamin addition. The provided vitamins were: (list). In this first experiment, the carbon sources were a mixture of 20 carbon substrates (glucose, myo-inositol, sorbitol, sucrose, xylose, 2-oxoglutarate, citrate, malate, pyruvate, succinate, GABA, glutamate, glycine, L-alanine, D-alanine, β-alanine, L-arginine, D-arginine, putrescine and urea). Each carbon source was provided at 36 mg C per L, such that the combined carbon concentration in the medium was again 720 mg C per L. These diverse carbon substrates were provided in order to maximise the probability of observing a positive growth phenotype, unobscured by substrate preference effects. Next, we investigated the specific vitamins required for the growth of strains Opi and Cpu, this time providing single carbon sources of either alanine (for Opi) or glucose (for Cpu). These substrates were chosen following preliminary experiments that identified these compounds as preferred single carbon substrates for each strain. Here, two approaches were used to pinpoint vitamin auxotrophies: either the addition of a single vitamin (V1 approach) or the removal of a single vitamin from the eight-vitamin mix (V7 approach). In both approaches, vitamins were provided at the previously used concentrations.

To integrate our experimental observations of vitamin auxotrophy with computational predictions of vitamin biosynthesis capacities, we interrogated the computational genome annotations for these genetically sequenced strains using the IMG database (22). These gene annotations were cross-referenced against a list of EC numbers for all characterised enzymes of bacterial vitamin synthesis, taken from Rodionov et al (23).

## Results

### Exometabolomic profiling of the SynCom on maize root extract

First, we hypothesised that the stable assembly of this community could be the result of metabolic resource partitioning, whereby the strains avoid direct competition by occupying differential metabolic niches in the maize rhizosphere habitat. To investigate this, we undertook a high-throughput LC-MS exometabolomic analysis to identify which maize root metabolites the strains consume as growth substrates. Methodologically, this involved cultivating the microbes on a growth medium where maize root extracts were the sole carbon source, then analysing the derived culture supernatants using LC-MS. The derived data enabled us to profile how the strains had modulated their chemical environment, providing new information about the maize root metabolites consumed by each strain, and the microbe-derived metabolites released into the growth medium.

Our data analysis strategy initially focussed on characterising microbial substrate preferences, by identifying the metabolite peaks that were present in the sterile medium but depleted following bacterial growth. The derived data is presented as a heatmap of 425 metabolite ions depleted by at least one strain (Fig 1a). Investigating the metabolite depletion profiles, we observe clear evidence of niche differentiation, because each strain depletes a distinct set of metabolite ions. Intriguingly, the C7 seems to combine the metabolic capabilities of each individual strain, with this ‘addition effect’ meaning that the C7 occupies a much broader metabolic niche compared to any single individual.

**Figure 1:**
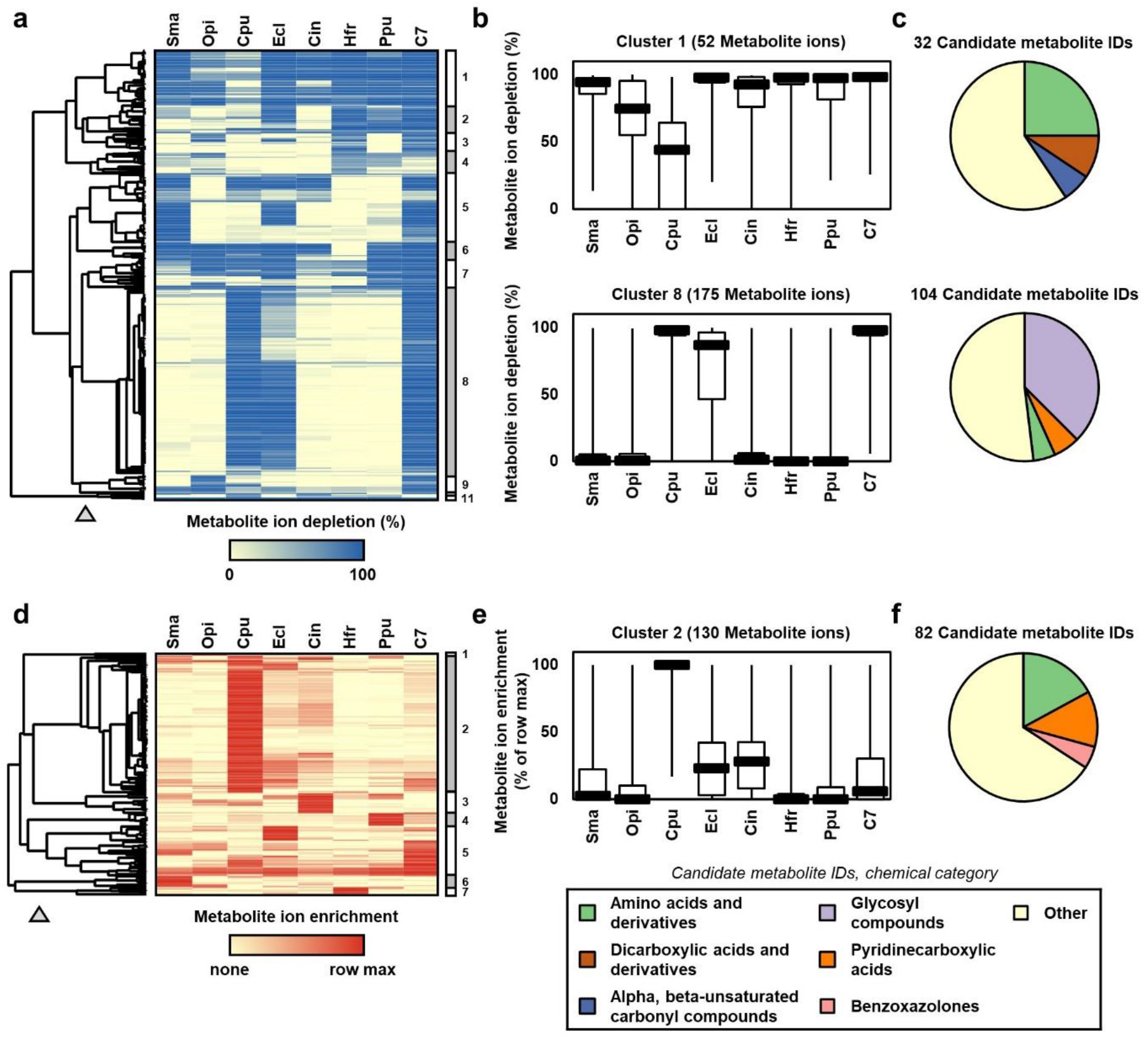
Metabolic footprints of the SynCom and constituent strains on maize root extract. (A) Heat map showing metabolite depletion profiles for 425 metabolite ions present in maize root extract, measured by LC-MS exometobolomics. Metabolite ions were included if they were depleted by at least one inoculum (LC-MS abundance<50% compared to sterile control, p<0.05), cell colour represents the mean depletion value from three independent replicates. Rows were clustered via Pearson correlation. The numbered panel to the right of the heatmap shows 11 clusters of metabolite ions, the grey triangle indicates where the dendogram was cut to split these clusters. (B) Box plots of metabolite ion depletion for two metabolite ion clusters in panel A. Cluster 1 contains metabolite ions generally depleted by all seven strains and the C7, whereas cluster 8 contains metabolite ions that were primarily depleted only by strains Cpu, Ecl and the C7. (C) Pie charts showing the chemical category of the candidate metabolite IDs for these two clusters. Candidate IDs were generated by searching the m/z value of the depleted metabolite ion against the Metlin database, and then the Classyfire database was used to categorise the chemical class of the top hit. (D) Heat map of metabolite enrichment profiles for 228 metabolite ions that exhibited higher abundance in the inoculated samples versus sterile controls (abundance>200%, p<0.05). Cell colour represents the mean enrichment value from three independent experiments, measured via untargeted LC-MS exometabolomics. Rows were clustered via Pearson correlation. The numbered panel to the right of the heatmap shows eight clusters of metabolite ions, the grey triangle indicates where the dendogram was cut to split these clusters. (E) Box plot of metabolite ion enrichment for cluster 2, which were primarily enriched only by strain Cpu. (F) Pie chart showing the chemical category of the candidate metabolite IDs for this cluster. Candidate IDs were generated by searching the m/z value of the depleted MS-feature against the Metlin database, and then the Classyfire database was used to categorise the chemical class of the top hit.

We sought to gain more information about the chemical identity of these depleted metabolite ions. There is considerable uncertainty in these data, because our workflow involved matching the m/z values of these high-resolution MS measurements against the Metlin database. Therefore, we name these ‘Candidate metabolite IDs’, which were then grouped into chemical category according to the Classyfire database. Our aim was not to generate a comprehensive list of all detected metabolites, but instead to narrow down which metabolites niches are occupied by the individual strains.

To present the data, we integrate the heatmap of metabolite depletion profile with the chemical categorisations of the candidate metabolite IDs. The heatmap dominated by a large grouping of depleted metabolite ions that were only depleted by three inocula: Ecl, Cpu and C7. Chemically, many of these m/z values were categorised in the group ‘glycosylated compounds’, and closer inspection of the data indicated that this cluster includes a large number of maize secondary metabolites, including sugar conjugates of flavonoids and benzoxaolones (Table S1). The overlapping niches of these two strains is notable, because the work of Niu et al (2017) found that the absence of Ecl allows Cpu to dominate the assembled community.

The heatmap also contains a relatively small cluster of 52 metabolite ions that were universally depleted by all seven strains. Chemical categorisation showed that this cluster contained a large proportion of primary metabolites, such as amino acids and organic acids. We validated this observation using GC-MS exometabolomic profiling of the same samples, which showed that the consumption of major sugars, organic acids and amino acids was commonly exhibited by all seven strains (Fig S1). Assessing this result, we conclude that primary metabolites probably play a minor role in mediating niche differentiation, although they clearly represent important growth substrates.

Our second data analysis strategy with the untargeted LC-MS exometabolomics dataset involved determining which microbe-derived metabolites were released into the growth medium by each strain. This heatmap is dominated by a large cluster of metabolite ions predominately released by strain Cpu (Fig 1d). Chemically, many of these m/z values were categorised as ‘amino acids and derivatives’, and include several free benzoxazolones lacking the sugar group (Fig 1e-f, Table S2). Due to Cpu’s potential to dominate the other strains, we suggest that metabolite ions could represent antimicrobial compounds or other chemical antagonists.

### Substrate utilisation profiles of the SynCom and C6 drop-outs

Although our exometabolomics workflow provided a high-throughput method to measure the metabolic footprints of the SynCom strains, the major disadvantage of this approach is that there is considerable uncertainty regarding the chemical identity of the detected metabolite ions. Therefore, we undertook an orthogonal methodology to assess metabolic niche occupancy of the SynCom strains, by measuring their substrate utilisation profiles using via BIOLOG EcoPlate. This well-established method measures the metabolic activity of microbes cultivated in a 96-well plate, wherein each well is pre-loaded with a single chemical substrate as the sole carbon source.

Our first experiment involved measuring the substrate utilisation profiles of all seven individual strains and the C7. The derived heatmap clearly discriminates the strains according to distinct metabolic niches, such as Ecl (carbohydrates), Cin (polymers) and Ppu (amines and amides) (Fig 2). This reaffirms our previous observation that the C7 combined the substrate utilisation capacities of all constituent strain, which again illustrates the ‘addition effect’ that was consistently observed in all three metabolic profiling methodologies used in this study (Fig S2).

**Figure 2:**
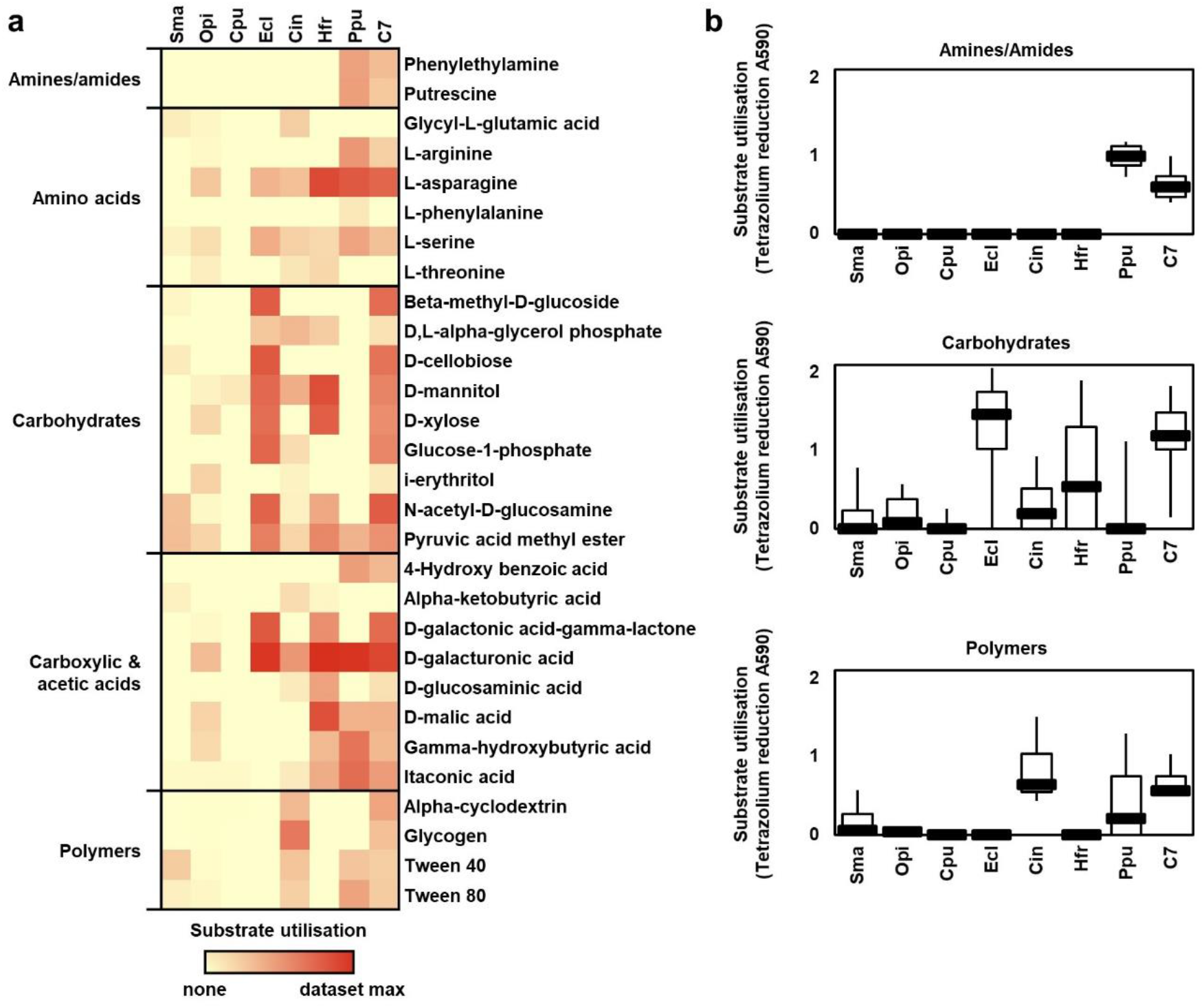
Metabolic niche profiling of the SynCom and constituent strains using phenotype microarray. (A) Heat map of substrate utilisation on 29 carbon sources by the SynCom strains measured using BIOLOG EcoPlates. Cell colour represents mean A590 value of three independent replicates. (B) Box plots of substrate utilisation for three selected chemical categories.

We undertook a second experiment, which assessed how much influence each strain exerts over the metabolic activity of the C7. Here, we undertook C6 drop-out experiments analogous to the approach of Niu et al (11), by removing individual strains from the community. Results showed that removal of strains Ecl and Ppu made the biggest impact on substrate utilisation. Specifically, removal of Ecl manifested in lower utilisation of carbohydrates, whereas Ppu removal resulted in lower utilisation of amino acids and nitrogenous compounds (Fig 3a). We plotted these data using PCA (Fig 3b), and then integrated our results with the study of Niu et al (11) by comparing their community assembly data versus our metabolic activity profiles (Fig 3c). There is clear concordance between the two datasets, with both studies showing that C6 communities lacking Ecl and Ppu were the two groups exhibiting the largest differences to C7, in terms of both community assembly and metabolic function. This correlative link provides evidence that metabolic properties can help to explain why certain strains exert a disproportionate influence on community assembly.

**Figure 3:**
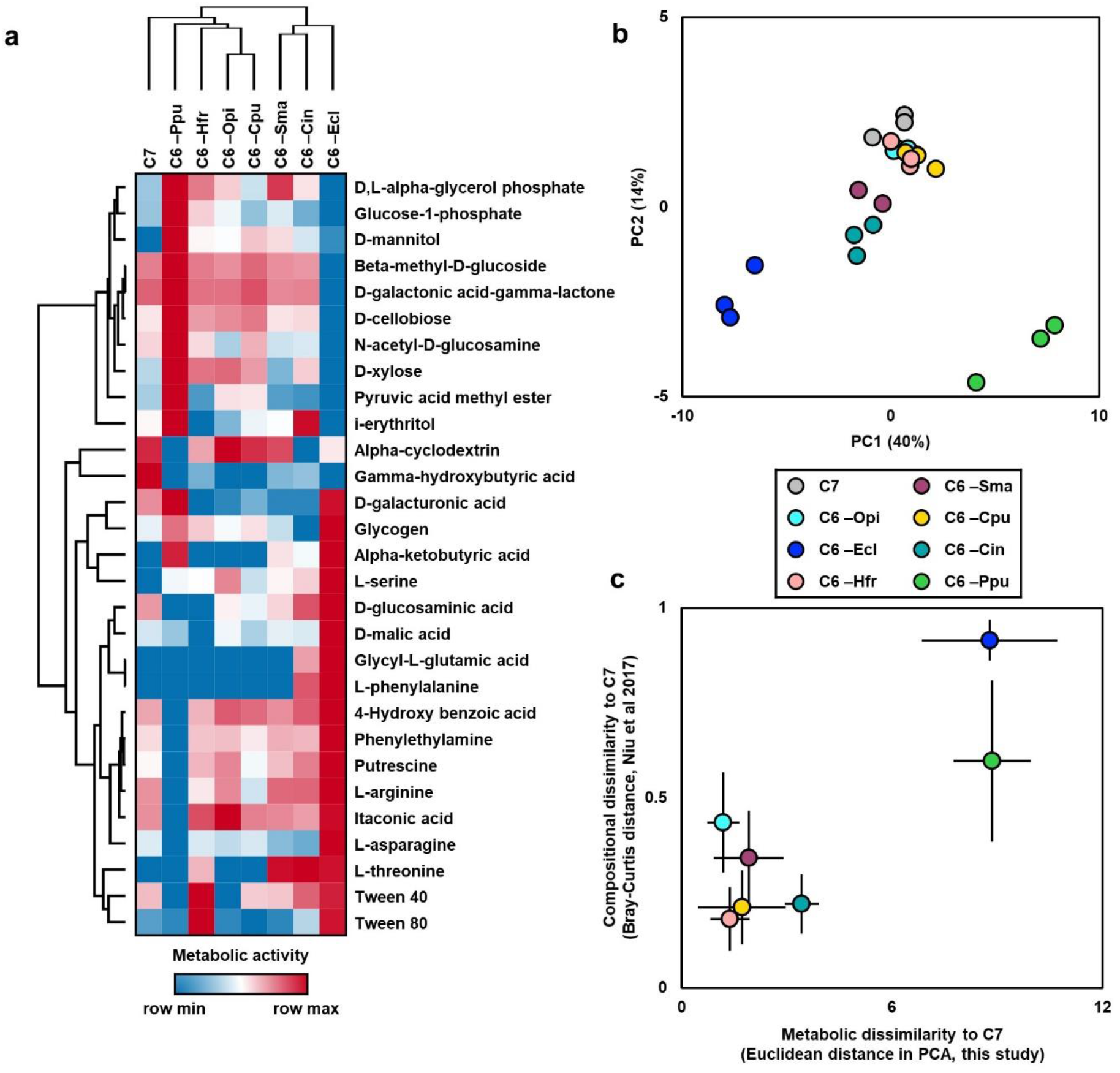
Metabolic analysis of C6 ‘drop-out’ communities via phenotype microarray. (A) Heat map of substrate utilisation for the C6 communities where one strain was removed, as well as the intact C7 SynCom, measured using BIOLOG EcoPlate. Cell colour represents mean value of three independent replicates. (B) PCA of metabolic phenotypes for the C6 communities and the C7. (C) Scatter plot comparing the compositional dissimilarity of C6 communities versus C7 on maize roots previously measured in Niu et al (2017, PNAS 114:E2450) versus the metabolic dissimilarity of phenotype microarray profiles versus C7 measured in this study (Fig 3B). Error bars represent SD.

### Cross-feeding phenotypes of the SynComs strains

We then explored pairwise cross-feeding interactions amongst SynCom strains. This initially involved determining which of the seven strains could grow on a minimal medium without any additional vitamins (Fig 4a). Across nine carbon substrates, only Ecl, Hfr and Ppu exhibited positive growth phenotypes, with Ecl generally growing fastest (Fig 4b).

**Figure 4:**
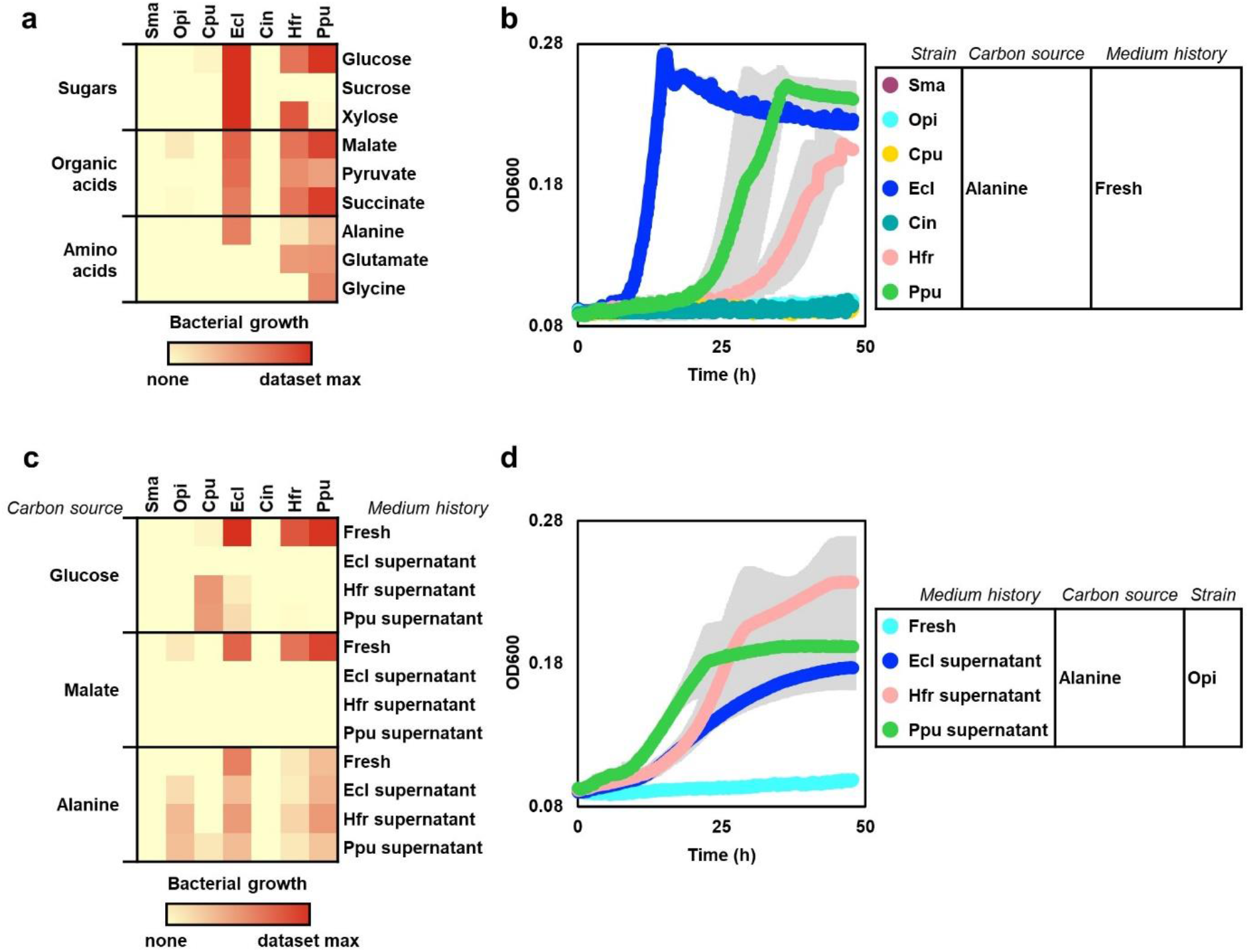
Growth assays and cross-feeding phenotypes of SynCom strains on minimal media. (A) Heat map of growth phenotypes for the individual SynCom strains cultivated on minimal media using nine sole carbon sources. Colour intensity corresponds to mean growth performance, measured via area under the curve in three independent experiments (B) Growth curves of all seven strains cultivated on minimal medium with alanine as sole carbon source. Data points represent the mean value of three independent replicates, grey shadings represent SD. (C) Heat map of cross-feeding phenotypes for the seven SynCom strains grown on culture supernatants harvested from either Ecl, Hfr and Ppu strains, following their pre-cultivation on either glucose, malate or alanine as sole carbon source. Colour intensity corresponds to mean growth performance, measured via area under the curve in four independent experiments. (D) Growth curves of the Opi strain cultivated on either a ‘fresh’ minimal medium with alanine as the sole carbon source, or on culture supernatants harvested from either Ecl, Hfr and Ppu, following their pre-cultivation on minimal medium with alanine as the sole carbon source. Data points represent the mean value of four independent replicates, grey shadings represent SD.

Once we had defined these three strains as prototrophic, our next step was to conduct cross-feeding assays, by harvesting filter-sterilised culture supernatants from the three prototrophic strains to provision as growth media for all seven strains (Fig 4c). This dataset revealed that the effectiveness of cross-feeding involves a complex interplay of effects relating to donor strain, recipient strain and carbon source. For instance, the organic acid malate did not support cross-feeding in any of the studied strains, whereas the amino acid alanine is relatively effective at supporting cross-feeding for five recipient strains. There is little evidence of ‘donor effect’, because spent media harvested from all three prototrophic strains (Ecl, Hfr and Ppu) have generally similar potential for nourishing the recipients. Analysing ‘recipient effects’ one observation of particular interest is the ‘growth boost’ received by Opi, which cannot grow on fresh alanine medium, but grows effectively on culture supernatant harvested from three other SynCom members (Fig 4d). Taken together, these data provide experimental evidence that cross-feeding can occur in this SynCom, and categorises the strains into three potential donors (Ecl, Hfr and Ppu) and two key recipients (Opi and Cpu).

#### B-vitamin dependencies of the SynCom strains

Literature evidence shows that B-vitamin auxotrophy is widespread amongst bacteria, with the exchange of B-vitamins positioned as a key mechanism for maintaining the diversity of microbial communities (9). Here we investigated the B-vitamin responses of all seven SynCom strains, by studying their growth curves either with or without the addition of an eight-vitamin mixture containing all B-vitamins, in media containing a diversity of simple carbon substrates (Fig 5a). Data indicate that the strains can be grouped into three general categories: 1) High-responders (Opi and Cpu) that receive a growth boost from B-vitamins; 2) Non-responders (Ecl, Hfr and Ppu) that exhibit strong growth regardless of vitamin addition, and 3) Unknown (Sma and Cin) that did not grow in either condition. To demonstrate the magnitude of this differential vitamin responses, we show raw growth curves for two exemplary strains (Opi and Ecl) (Fig 5b). This showed that Ecl grows almost identically under the two conditions, whereas Opi receives a dramatic growth benefit from exogenous vitamins.

**Figure 5:**
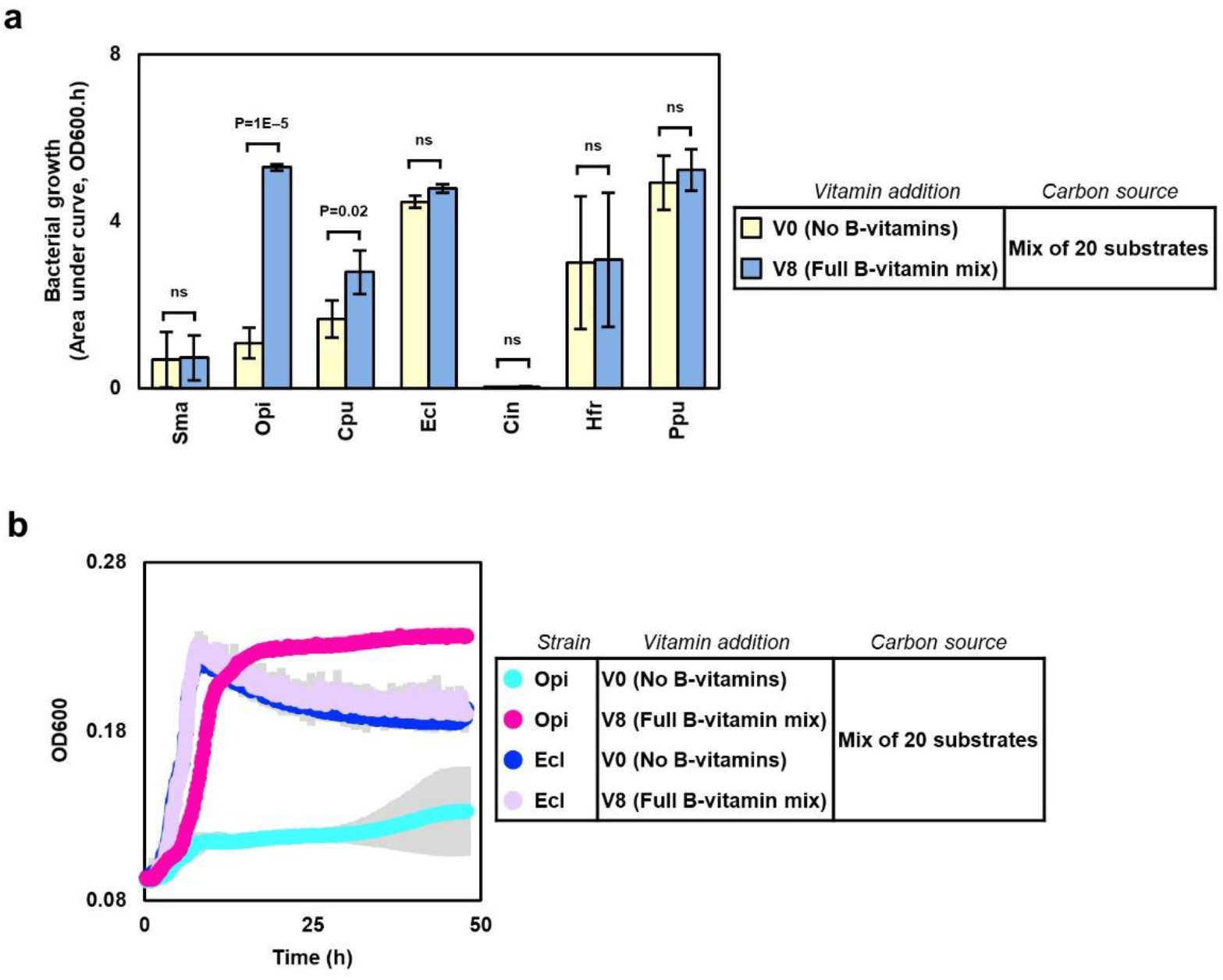
B-vitamin responses of the SynCom strains. (A) Bar chart showing the growth performance of each strain either with or without B-vitamin mixture. Statistical significance of each strain’s vitamin response was determined using Student’s t-test. Error bars represent SD, n=4. (B) Growth curves of strains Opi and Ecl either with or without B-vitamin mixture. Data points represent the mean value of four replicates, grey shadings represent SD.

Overall, this profile of vitamin dependency is largely consistent with the results of cross feeding assays (Fig 4c). Across the two datasets, the same strains which were grouped as ‘cross-feeding donors’ (Ecl, Hfr and Ppu) show no requirement for exogenous vitamins, whereas the strains that were grouped as ‘key recipients’ (Opi and Cpu) receive a growth benefit from the addition of a vitamin mixture. This provides indirect evidence that vitamin exchange between prototrophs and auxotrophs could play a role in mediating stability of this SynCom.

Our next step was to undertake reductionist experiments to pinpoint which particular vitamins were required by the auxotrophic strain Opi. We first hypothesised that Opi was auxotrophic for one specific B-vitamin, but this was proven false, because the provision of any single vitamin failed to rescue Opi’s growth phenotype (Fig 6a). Our next step was to undertake V7 ‘drop-out’ experiments, by removing one single vitamin from the 8-vitamin mixture. Results of this experiment revealed that a significant drop in Opi’s growth performance was elicited by two of the V7 mixes (V7 -Thiamine and V7 -Biotin) (Fig 6b). This provided negative evidence that Opi is a double-auxotroph for thiamine and biotin, and we next corroborated this finding with positive evidence, showing the addition of these two vitamins rescues Opi’s growth phenotype in a manner similar to the V8 (Fig 6c). Taken together, we conclude that Opi is auxotrophic for both thiamine and biotin, which therefore positions the exchange of these metabolites as one candidate mechanism mediating the stability of this SynCom.

**Figure 6:**
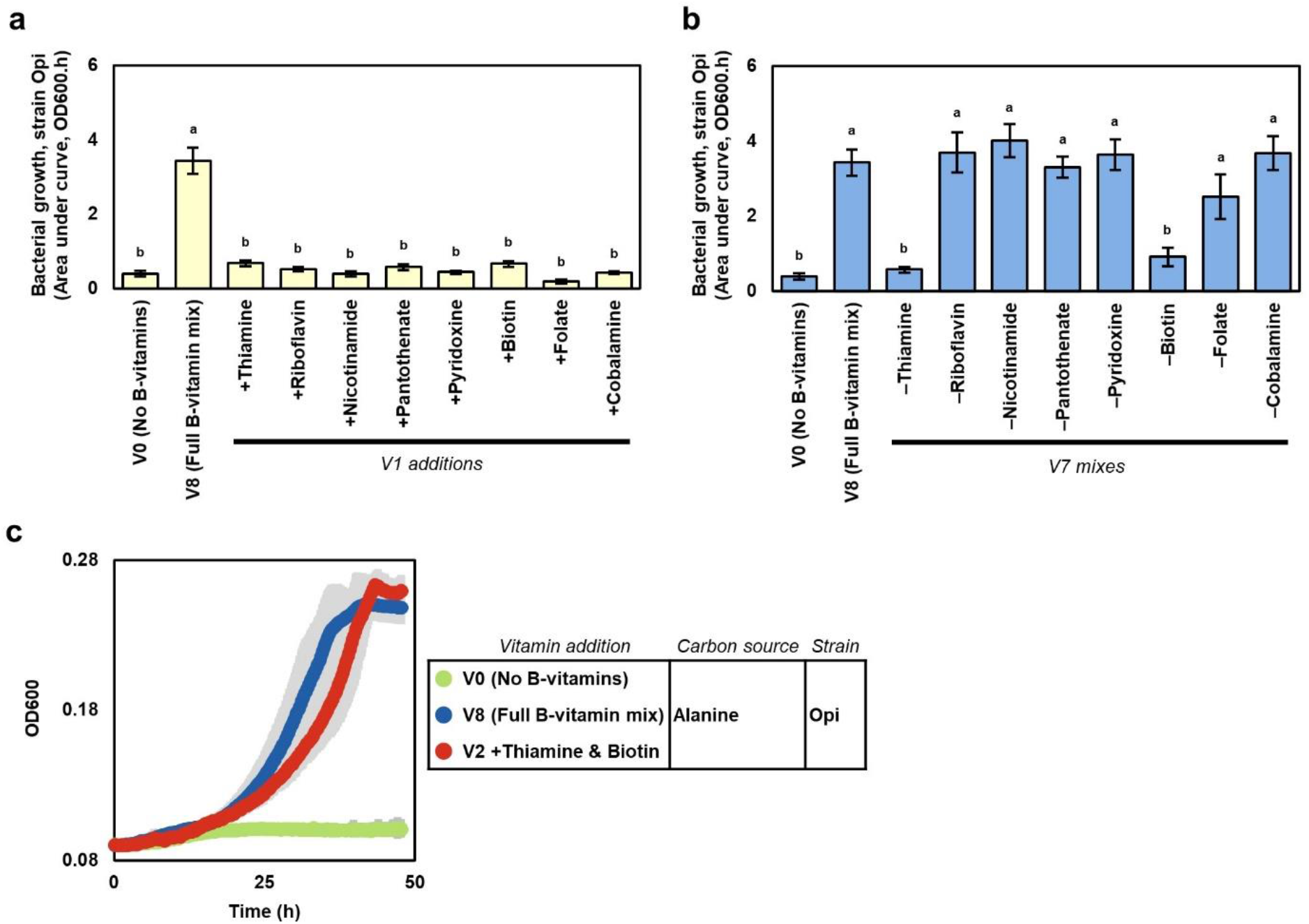
Pinpointing B-vitamins required for growth of strain Opi. (A) Bar chart showing the effect of adding single B-vitamins on Opi growth performance. Groups annotated with the same letter are not significantly different using Tukey’s HSD test (α=0.95). Error bars represent SD, n=4. (B) Bar chart showing the how Opi’s growth performance is affected by V7 mixes, where on single B-vitamin was removed from the eight-vitamin mixture. Groups annotated with the same letter are not significantly different using Tukey’s HSD test (α=0.95). Error bars represent SD, n=4. (C) Growth curves of Opi strain cultivated either with no B-vitamins, with a full mixture of eight B-vitamins, or with a V2 addition of Thiamine and Biotin. Data points represent mean value of four replicates, grey shadings represent SD.

We also investigated the specific vitamins required for the growth of strain Cpu. This revealed that Cpu is auxotrophic for seven out of eight B-vitamins (all except for B12, cobalamine) (Fig S3). This suggests that Cpu has a vitamin-scavenging lifestyle, which is intriguing considering that this strain has the potential to dominate the community assembly in the absence of Ecl (11), and also that it releases a much larger set of metabolites compared to any of the other strains (Fig 1d).

Our next step was to assess how accurately vitamin auxotrophies can be predicted by genome annotation. All of these strains were previously genetically sequenced (24), and we used IMG’s computational gene annotations of these sequence data to predict the vitamin biosynthesis capacity of these strains, and then compared these predictions to our experimental measurements. Results show that vitamin auxotrophy is correctly predicted in 77% of cases (27/35) (Fig S4). This suggests that computational approaches might play a useful role in preliminary screening of auxotrophic strains, but that genetic data in isolation does not yet have the capacity to fully predict vitamin auxotrophies, with laboratory experiments being necessary to provide a full characterisation of bacterial vitamin requirements.

## Discussion

The motivation of this study was to define specific metabolic mechanisms that mediate strain coexistence in an established bacterial SynCom, by exploring the processes of niche differentiation and cross-feeding. Using multiple methods, we comprehensively documented the metabolic niches occupied by each strain, which demonstrated that the SynCom members exhibit metabolic complementarity. Furthermore, we described which specific strains are the donors and recipients of cross-feeding interactions, and showed that some strains were auxotrophic whereas others are prototrophic. Results of vitamin dependency experiments implicated certain B-vitamins as candidate molecules exchanged between strains. Interpreting our results, we are particularly interested in defining metabolic traits which could serve as ‘design principles’ for rationally assembling stable SynComs.

### Metabolic complementarity as a mechanism underpinning SynCom stabilty

The ecological principle of niche differentiation states that species which coexist in the same habitat must consume different resources in order to avoid competition (25). This principle could guide efforts to compose stable microbial communities for applications in agriculture and medicine, by matching compatible strains according to their non-overlapping substrate preferences (1). The SynCom investigated in this study was previously shown to assemble stably on maize roots (11), and therefore, we postulated that the constituent strains would occupy complementary metabolic niches.

Using multiple experimental approaches, we conclusively demonstrate that the strains in this SynCom exhibit metabolic niche complementarity, This is shown by three orthogonal lines of evidence: 1) Metabolomic footprinting on maize root extracts, which pinpoint hundreds of metabolites only consumed by individual strains, tentatively matched to maize secondary metabolites; 2) Phenotype microarray results that the strains utilise different classes of metabolic substrates (ie: polymers, carbohydrates, amides); and 3) Vitamin auxotrophy profiling, which clearly distinguished between a set of three prototrophic strains versus two auxotrophs. Interpreting these findings, we postulate that metabolic complementarity could be a causative mechanism mediating stability in this SynCom. Potentially, future experiments could test this principle across a larger number of differentially formulated SynComs. For example, would SynComs composed from strains with overlapping substrate preferences exhibit a collapse in diversity due to competitive exclusion effects? In turn, can SynCom stability be promoted by combining strains that occupy non-overlapping metabolic niches? And would it be possible to engineer interdependence between strains, by combining vitamin exporters with reciprocal auxotrophs?

In the literature, the primary rationale guiding SynCom assembly has been taxonomic representativeness, whereby individual SynCom strains are selected as representative members corresponding to the major phylogenetic groupings which occur in naturally assembled microbiomes (26). It is often presumed that taxonomically diverse communities will inevitably exhibit functional diversity, because the different microbial phyla have committed to diverging ecological strategies early in their evolutionary history (27,28). However, there is increasing evidence that bacterial metabolic traits are poorly predicted by taxonomy, with closely related strains often showing widely diverging substrate preferences, potentially mediated the horizontal transfer of metabolic genes (29). In our opinion, the concept of metabolic complementarity could be a useful tool for designing stable SynComs, because it has a stronger mechanistic basis for promoting stability compared to the standard approach of combining diverse taxa.

### New insights into the metabolic properties of a keystone strain

Keystone strains are of major interest in the field of microbiome engineering, because the presence of these strains promotes the diversity of the surrounding community. Therefore, keystone strains could potentially be administered in cases of microbiome dysbiosis to restore community diversity, or alternatively included into SynComs to promote the coexistence of accompanying strains (30). There is significant interest in defining which members of the plant microbiota are keystones (31), and the work of Niu et al (11) provided a useful resource to the field, because it empirically defined Ecl as a keystone strain stabilising SynCom assembly on maize roots. Here, our results build upon this previous work, by providing new mechanistic information about the metabolic properties of Ecl in relation to the other SynCom strains and also the C7.

In this study we show that Ecl is a fast-growing strain with a broad metabolic niche and a large metabolic influence on the C7. It can rapidly utilise a diverse range of carbon substrates without the need for exogenous vitamins, and its culture supernatants can sustain other strains in cross-feeding assays. We postulate that these results provide a mechanistic explanation for the keystone behaviour of Ecl demonstrated in Niu et al (11). Specifically, one element of Ecl’s keystone behaviour could be its overlapping metabolic niche with the disruptive Cpu (Fig 1). Potentially, the faster-growing Ecl could outcompete Cpu for the primary usage of these maize secondary metabolites, which could prevent Cpu from dominating the community because these plant-substrates could serve as building blocks for antimicrobial compounds synthesised by Cpu. A further element could be Ecl’s cross-feeding ability, whereby this prototrophic strain synthesises vitamins that are subsequently released into the environment to promote the coexistence of auxotrophic strains. Future experiments could test whether the metabolic properties exhibited by Ecl are general characteristics shared by all keystone strains, or whether they are only relevant in the context of this simplified SynCom. If these metabolic characteristics are indeed common amongst keystones, then potentially they could be used as criteria for identifying candidate keystone strains from large panels of microbial isolates.

### Tailoring stable SynComs for specific plant genotypes

Plant species differ in their root chemistry (32), and there is growing evidence that species-specific secondary metabolites act as a selection pressure shaping the composition of the rhizosphere microbial community (33). This has an impact on the design of plant-associated SynComs, because microbial strains need to be equipped to colonise the chemical environment unique to the target host plant. The Niu et al (11) SynCom was assembled via host-mediated selection on maize roots, which automatically means that these strains are competent in the maize root environment. In this study, we provide new information about the metabolic niches occupied by each strain, using LC-MS exometabolomics to infer that maize secondary metabolites represent a major source of differential metabolic niches, particularly for the strains Ecl and Cpu. However, we have not authenticated the identity of these maize root metabolites using reference standards, so further investigations are necessary before we can unequivocally identify the specific metabolites that nourish each strain. But nevertheless, we feel that the exometabolomic approach utilised in this study could be a useful tool in tailoring SynCom design for distinct plant species, because it could facilitate the optimal matching between host root chemistry versus microbial substrate utilisation.

### Conclusion

This study builds upon the work of Niu et al (11), by characterising the metabolic phenotypes exhibited by each member of their previously developed seven-strain bacterial SynCom which assembles stably on maize roots. We define the metabolic interplay between the strains, by documenting their differential metabolic niche occupancies and exploring donor-recipient cross-feeding interactions. We postulate that these metabolic mechanisms could be illustrative of general principles underpinning the stability of microbial communities, which could be used to guide the rational assembly of stable SynComs.

## Acknowledgements

We thank Ben Niu (Northeast Forestry University, China) for providing bacterial strains, Nicole Mantke (Institute of Geology and Mineralogy, University of Cologne) for performing TOC measurements, and the Service Unit for Metabolomics and Lipidomics at the University of Göttingen for GC-MS access. Research at CEPLAS is funded by the Deutsche Forschungsgemeinschaft (DFG) under Germany’s Excellence Strategy – EXC 2048/1 – project 390686111. TI is funded by a Heisenberg Grant from the DFG (IS 273/10). RPJ was supported by a Humboldt Research Fellowship.

## Competing Interests

The authors have no competing interests.

## Supplementary Figure Legends

**Supplementary Figure S1:**
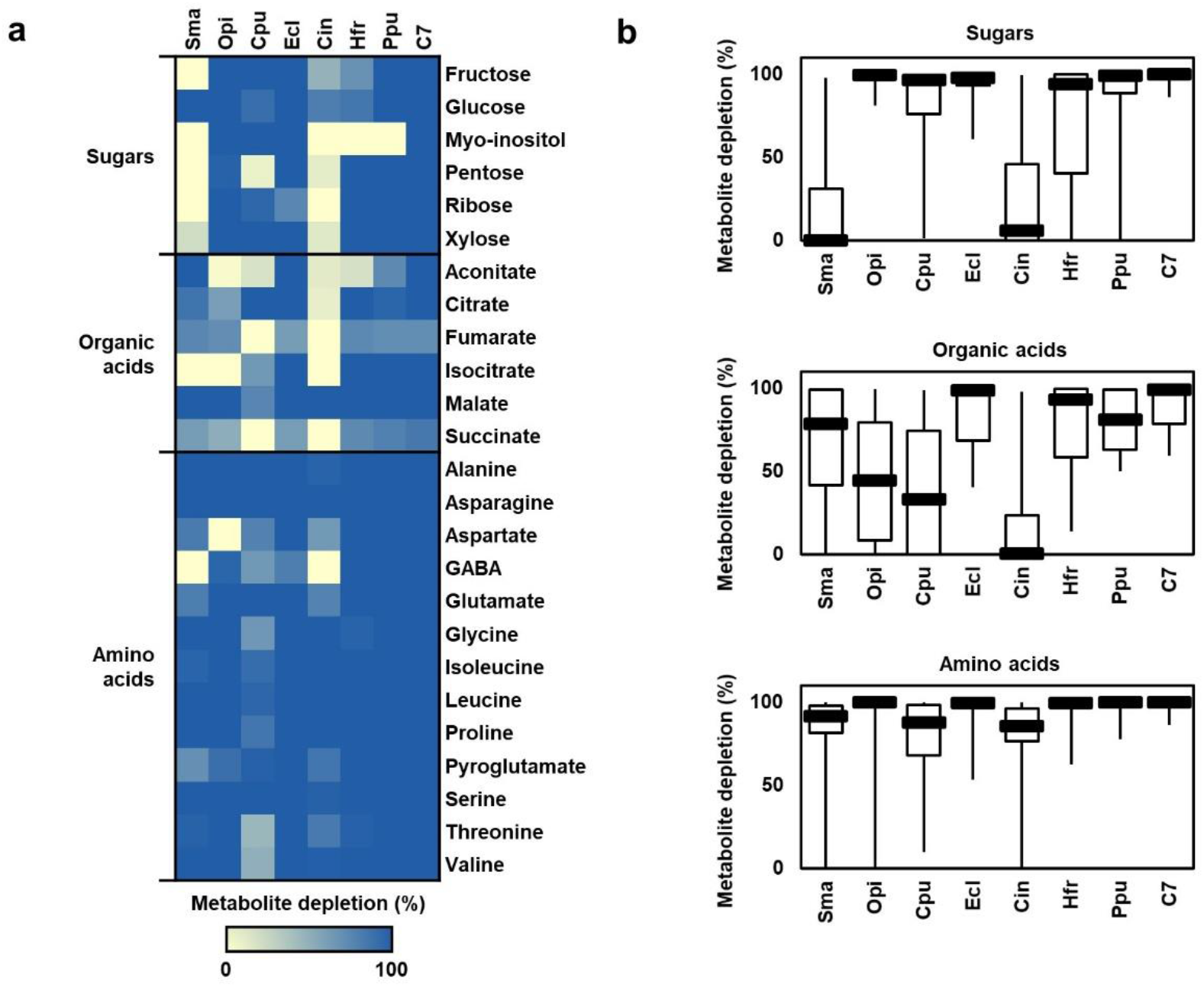
Depletion of primary metabolites from maize root extract by the SynCom and constituent strains. (A) Heat map of primary metabolites depletion by the SynCom strains measured via GC-MS exometabolomics of SynCom strains cultivated on maize root extract. Cell colour represents mean metabolite depletion value of three independent replicates. (B) Box plots of metabolite depletion for three categories of primary metabolites.

**Supplementary Figure S2:**
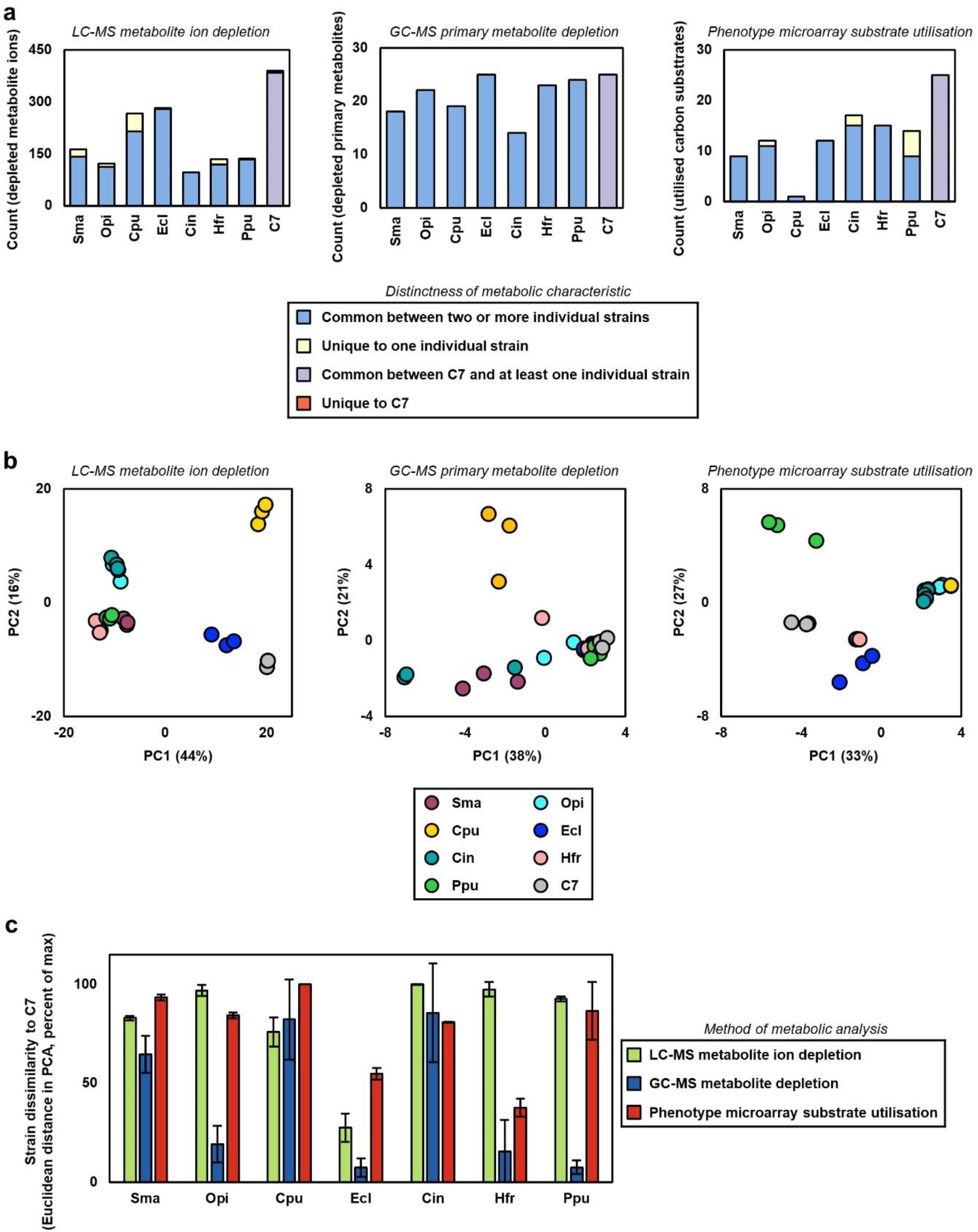
Overview of metabolic profiles for the SynCom and constituent strains measured via three methodologies. (A) Bar charts showing number of metabolite ions depleted by strains measured using LC-MS exometabolomics (metabolite ion abundance<50%, p-value<0.05), the number primary metabolites depleted by strains measured via GC-MS exometabolomics (metabolite abundance<50%, p-value<0.05), and the number of substrates utilised for growth by phenotype microarray measured via BIOLOG EcoPlate (A595>0.1, p-value<0.05). Bars are divided into common and unique metabolic characteristics. For the individual strains, this involved comparing that strain’s metabolic profile versus the six other individual strains (but not the C7), whereas for the C7, this involved comparing the C7 profile versus the seven individual strains. (B) PCAs of metabolic phenotypes measured via LC-MS exometabolomics (untargeted), GC-MS exometabolomics (targeted) and phenotype microarray (BIOLOG EcoPlate). (C) Bar chart of Euclidean distance between each individual strain versus the C7 SynCom for the PCAs in panel B. Here, smaller Euclidean distances indicate closer similarity between the metabolic profile of that individual strain versus the C7.

**Supplementary Figure S3:**
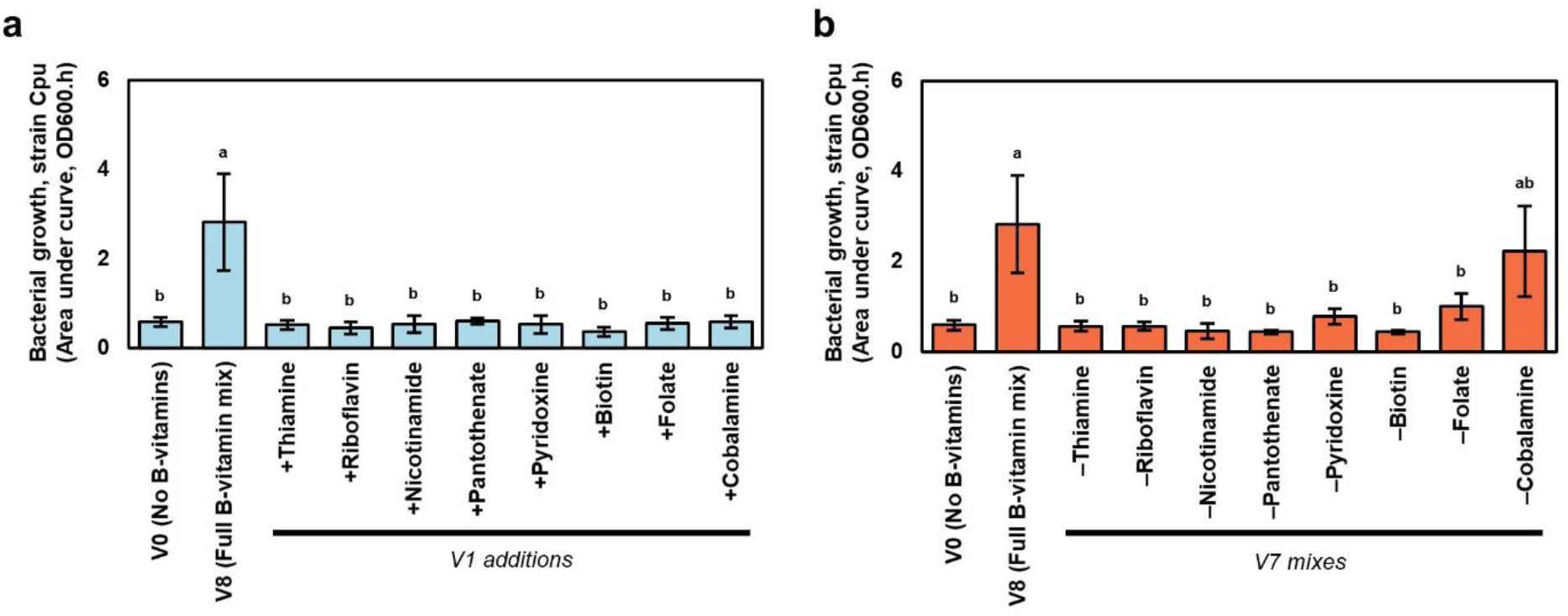
Pinpointing B-vitamins required for growth of strain Cpu. (A) Bar chart showing the effect of adding single B-vitamins on Cpu growth performance. Groups annotated with the same letter are not significantly different using Tukey’s HSD test (α=0.95). Error bars represent SD, n=4. (B) Bar chart showing the how Cpu growth performance is affected by V7 mixes, created by removing single B-vitamins from the eight-vitamin mixture. Groups annotated with the same letter are not significantly different using Tukey’s HSD test (α=0.95). Error bars represent SD, n=4.

**Supplementary Figure S4:**
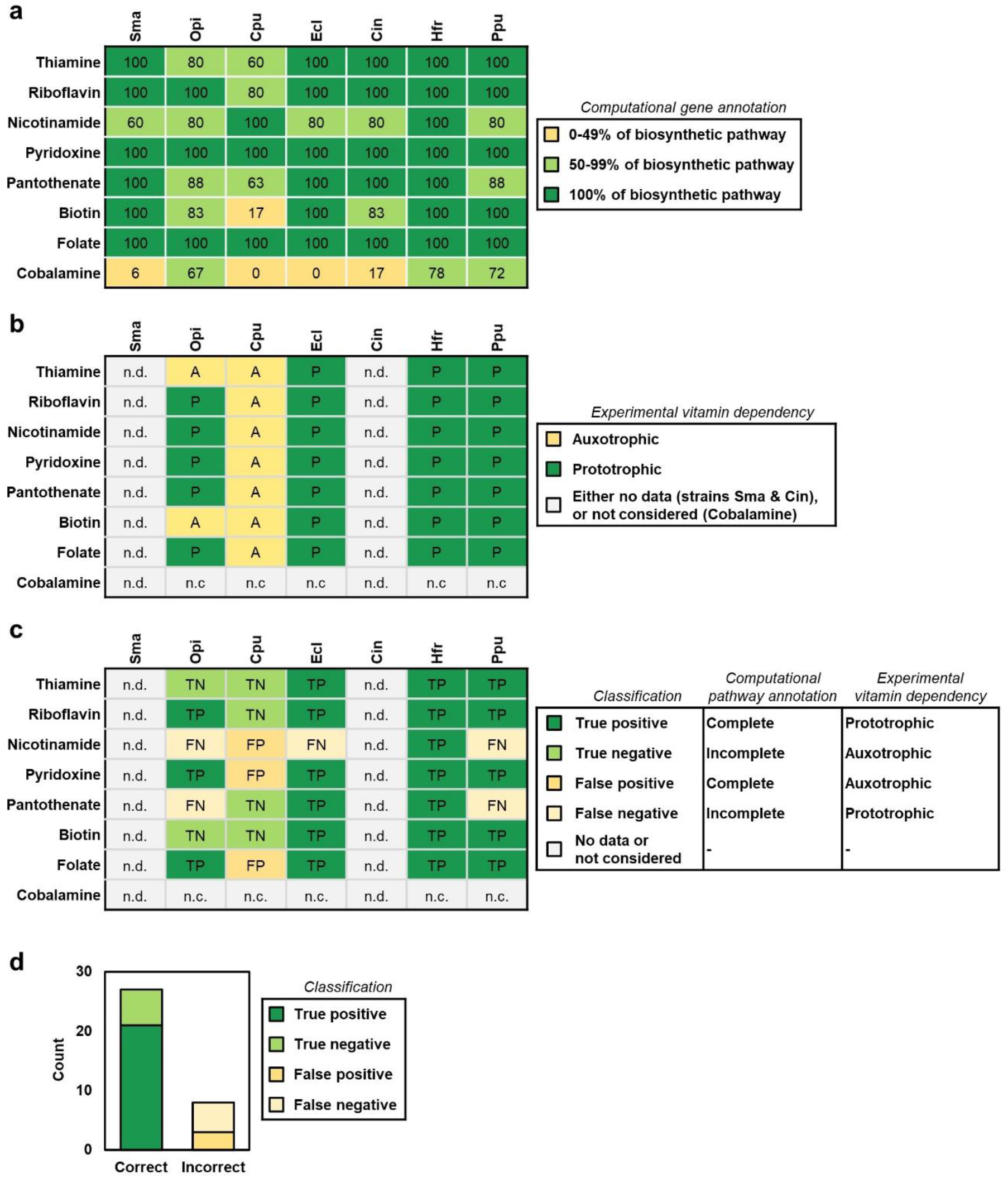
Concordance between computational prediction of vitamin auxotrophy versus experimental measurement of vitamin dependency for SynCom strains. (A) Table showing computational prediction of biosynthetic pathway completeness for eight B-vitamins in the seven strains. Table represents percentage of enzymatic steps in each vitamin biosynthetic pathway that are annotated as present in that strain’s genome, using IMG pipelines. Biosynthetic pathway enzymes follow Rodionov et al (2019, Front. Microbiol., 10:1316) (B) Table showing experimental B-vitamin dependency measured for the seven SynCom strains. Phenotypes of prototrophy and auxotrophy are inferred from the experimental data reported in Fig 5, Fig 6, and Fig S3. Strains Sma and Cin were not considered for this analysis, because they did not exhibit growth on any of the minimal media used in this study. Furthermore, cobalamine (B12) was also not considered for this analysis, because experimental data indicate that none of these strains receive a growth benefit from exogenous B12, while computational analyses indicate that all strains’ genomes encode B12-independent enzymes for methionine synthesis and nucleotide metabolism, which together suggests that B12 is not an obligate nutrient for these strains. (C) Table showing concordance between computational predictions of vitamin auxotrophy versus experimental measurements of vitamin dependency. Entries were generated by integrating the predictions in panel A versus the experimental measurements in panel B (not considering strains Sma and Cin, or vitamin cobalamine). (D) Bar chart summarising the accuracy of computational predictions of vitamin auxotrophy for SynCom strains. Chart was generated by plotting the contents of panel C for the 35 considered entries (ie: seven vitamins in five strains).

## Supplementary Tables

***Table S1:*** Abundance values and LC-MS information for 425 metabolite ions present in maize root extract and depleted by at least one bacterial inoculum in LC-MS exometabolomic experiments.

***Table S2:*** Abundance values and LC-MS information for 228 metabolite ions enriched into the extracellular medium by at least one bacterial inoculum in LC-MS exometabolomic experiments.

***Table S3:*** Abundance values for 25 primary metabolites present in maize root extract and depleted by bacterial inocula in GC-MS exometabolomic experiments.

***Table S4:*** Substrate utilisation measurements for 29 carbon sources present on BIOLOG Ecoplate phenotype microarrays.

## Notes

### Competing Interest Statement

The authors have declared no competing interest.

